# Subcellular localization of truncated MAGEL2 proteins: insight into the molecular pathology of Schaaf-Yang syndrome

**DOI:** 10.1101/2024.01.22.576607

**Authors:** Mónica Centeno-Pla, Estefanía Alcaide-Consuegra, Sophie Gibson, Aina Prat-Planas, Juan Diego Gutiérrez-Ávila, Daniel Grinberg, Roser Urreizti, Raquel Rabionet, Susanna Balcells

**Affiliations:** Department of Genetics, Microbiology and Statistics, Faculty of Biology, University of Barcelona, IBUB, IRSJD, 08028, Barcelona, Spain; Centro de Investigación Biomédica en Red de Enfermedades Raras (CIBERER) - Instituto de Salud Carlos III, Spain; Clinical Biochemistry Department, Hospital Sant Joan de Déu, CIBERER, 08950, Esplugues de Llobregat, Spain

**Keywords:** clinical genetics, Schaaf-yang syndrome, nervous system diseases, MAGEL2 mutations, neurodevelopmental disorders

## Abstract

Schaaf-Yang syndrome (SYS) is an ultra-rare neurodevelopmental disorder caused by truncating mutations in *MAGEL2*. Heterologous expression of wild-type (WT) or a truncated (p.Gln638*) C-terminal HA-tagged MAGEL2 revealed a shift from a primarily cytoplasmic to a more nuclear localization for the truncated protein variant. We now extend this analysis to six additional SYS mutations on a N-terminal FLAG-tagged MAGEL2. Our results replicate and extend our previous findings, showing that all the truncated MAGEL2 proteins consistently display a predominant nuclear localization, irrespective of the C-terminal or N-terminal position and the chemistry of the tag. The variants associated with arthrogryposis multiplex congenita (AMC) display a more pronounced nuclear retention phenotype, suggesting a correlation between clinical severity and the degree of nuclear mis-localization. These results point to a neomorphic effect of truncated MAGEL2, which might contribute to the pathogenesis of SYS.

## INTRODUCTION

*MAGEL2* (OMIM *605283) is one of the maternally imprinted protein-coding genes of the Prader-Willi region (15q11-q13). It is a single-exon gene, encoding one of the largest proteins of the type II MAGE protein family, comprising 1,249 amino acids. Within the N-terminal region of MAGEL2, there is a proline-rich domain that has been associated with RNA metabolism (1), while in the C-terminal end, spanning amino acids 1,027 to 1,195, lies a MAGE Homology Domain (MHD). MHD is a highly conserved sequence common to both type I and type II MAGEs that plays a crucial role in facilitating protein-protein interactions in retrograde endosomal transport or circadian rhythm processes (2,3). Since the identification of *MAGEL2* truncating mutations as the cause of Schaaf-Yang syndrome (SYS, OMIM # 615547) in 2013 (4), over 80 different mutations, all resulting in the production of truncated proteins lacking the MHD, have been reported.

SYS is an ultra-rare neurodevelopmental syndrome which shares some clinical features (neonatal hypotonia, developmental delay or sleep disorders) with the more common Prader-Willi syndrome (PWS, OMIM # 176270), caused by the lack of expression of paternal genes in 15q11-q13. However, individuals with SYS more frequently exhibit severe intellectual disability, autism spectrum disorder behaviors, and joint contractures (5). In addition, three particular *MAGEL2* truncating variants, p.Gln666Serfs*36, p.Leu708Trpfs*7, and p.Val991*, are associated with a significantly more severe phenotype characterized by arthrogryposis multiplex congenita (AMC), which ultimately leads to perinatal death (6–9).

In a previous study, to characterize the functional consequences of MAGEL2 truncation, we transfected vectors encoding hemagglutinin (HA)-tagged wild-type (WT) or one particular truncated MAGEL2 (MAGEL2-Gln638*) and assessed their subcellular localization (10). Immunocytochemistry (ICC) assays showed that in cells transfected with the WT construct, MAGEL2 was mainly located in the cytoplasm, while there was a shift towards the nucleus when transfecting the truncated protein. Importantly, this phenomenon was seen consistently across different recipient cell lines. However, it could be argued that the phenomenon was mutation-specific and or that the C-terminal HA tag might be mediating this effect.

To establish the generalizability of the results, we have now investigated a broader range of *MAGEL2* truncating variants. We generated additional plasmids expressing a N-terminal FLAG-tagged MAGEL2 protein, in which we introduced six truncating *MAGEL2* mutations as illustrated in Supplementary Figure 1. Four of them [p.Trp617* (c.1850G>A), p.Gln638* (c.1912C>T), p.Gln666Profs*47 (c.1996dupC), and p.Trp686Alafs*23 (c.2056_2066del)] were identified in SYS patients, while the other two [p.Gln666Serfs*36 (c.1996delC) and p.Leu708Trpfs*7 (c.2118delT)] have been associated with severe AMC and perinatal death (6–9).

## MATERIALS AND METHODS

ICC assays were conducted in transfected HeLa cells to detect the presence of FLAG and 4’,6-diamidino-2-phenylindole (DAPI). Cloning, cell culture and immunocytochemistry are described in Supplementary materials and methods. Images were acquired using a Zeiss confocal microscope LSM 880 and analyzed with ImageJ (11). To quantitatively assess the nuclear localization, we used Mander’s colocalization coefficient between the FLAG fluorescent signal and DAPI. Statistical significance was determined using one-way analysis of variance (ANOVA), and p-values less than 0.01 were considered significant.

## RESULTS

Our ICC assays revealed that all six heterologously expressed truncated proteins are predominantly found inside the nucleus, in contrast with the cytoplasmic localization of WT MAGEL2 (Figure 1). These findings were consistently observed across multiple experimental replicates, suggesting a genuine shift in the localization pattern induced by MAGEL2 truncating mutations.

**Figure 1.**
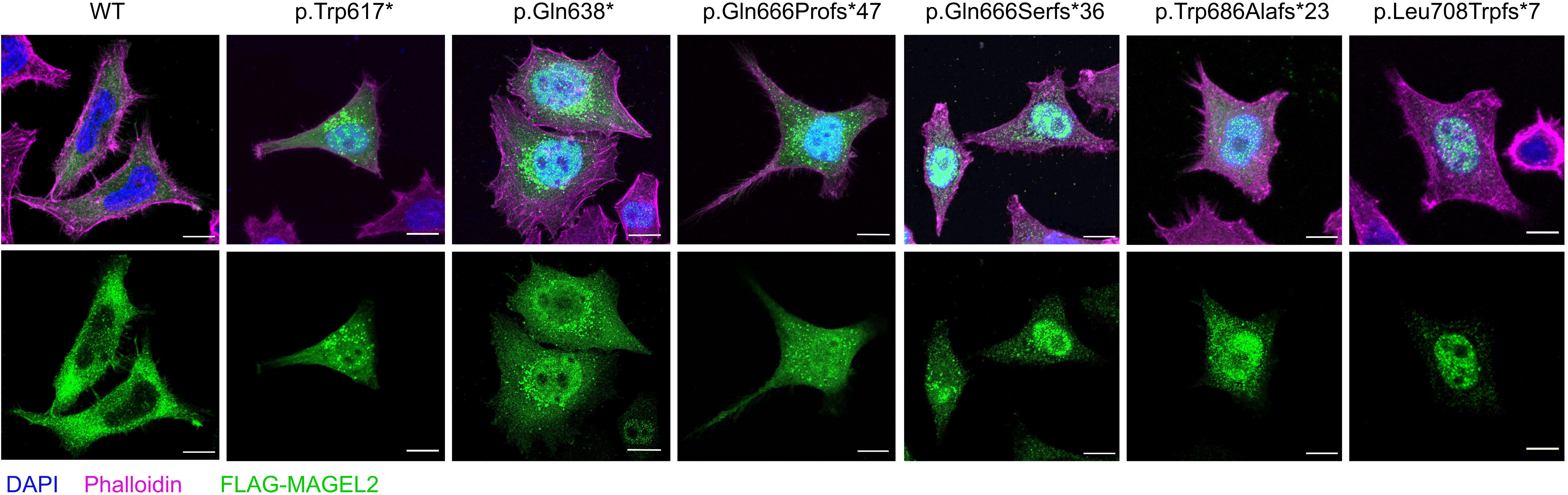
Representative immunofluorescence images of HeLa cells transfected with WT or truncated *MAGEL2* vectors carrying mutations p.Trp617*, p.Gln638*, p.Gln666Profs*47, p.Gln666Serfs*36, p.Trp686Alafs*23, and p.Leu708Trpfs*7 respectively, stained with DAPI (blue), Phalloidin (magenta), and anti-FLAG (green). Scale bar represents 10 µm.

A quantitative analysis of colocalization of MAGEL2 and DAPI revealed significant differences between all six truncated MAGEL2 variants and the full-length wild-type form and hinted at a connection between the degree of nuclear localization and the clinical severity of certain mutations (Figure 2). Notably, mutations p.Gln666Serfs*36 and p.Leu708Trpfs*7, which have been associated with severe AMC and perinatal death, exhibited a more pronounced nuclear localization. This finding adds a layer of complexity to our understanding of MAGEL2’s role in cellular processes, linking the subcellular behavior of its truncating variants to their phenotypic manifestation.

**Figure 2.**
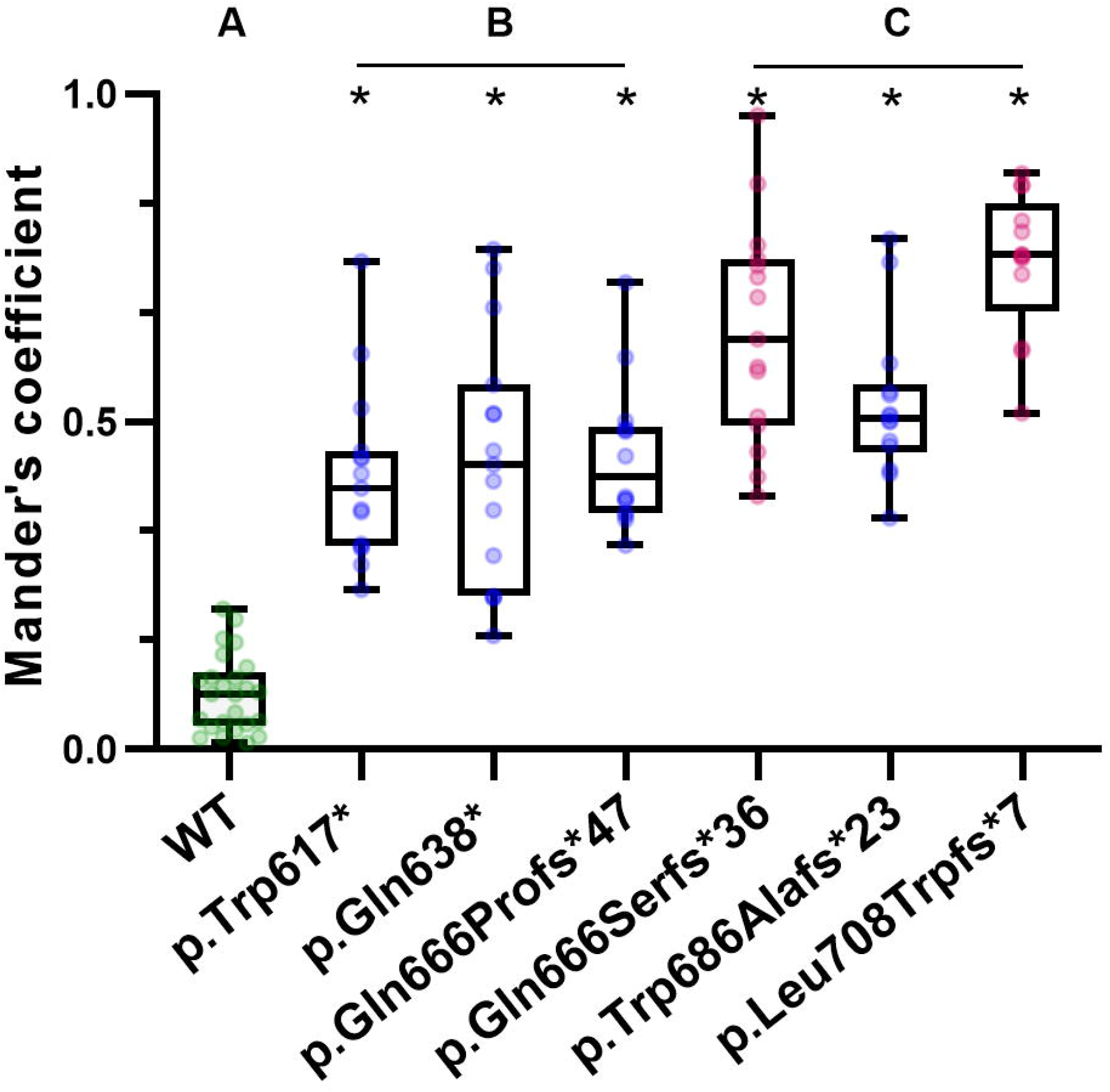
Mander’s coefficient quantification of the colocalization between the FLAG fluorescence signal and DAPI in cells transfected with WT FLAG-MAGEL2 or truncated FLAG-MAGEL2. Statistical analyses were performed using one-way ANOVA in GraphPad Prism. The symbol ‘*’ denotes significant differences of truncating mutations when compared to the WT group *p<0.01. Letters A, B, and C represent significant distinctions among groups identified through Tukey’s multiple comparisons. Data are presented as mean ± standard deviation from at least two independent experiments (N≥2).

## DISCUSSION

In our study, we made concerted efforts to address potential subcellular localization effects attributable to the tagging method. Our data consistently demonstrated a notable nuclear localization pattern of all exogenously expressed truncated MAGEL2 proteins. This strengthens the validity and specificity of our findings, suggesting that the observed localization reflects the inherent properties of truncated MAGEL2.

While the lack of a properly functioning antibody against the N-terminus of MAGEL2 hinders the validation of these findings with endogenous MAGEL2, the consistent and significant shift in localization points to a pathogenic neomorphic effect of the truncated protein in the nucleus. Furthermore, our findings suggest that the extent of nuclear localization of truncated MAGEL2 may play a role in the manifestation and severity of joint contractures in SYS patients. This correlation could be invaluable for predicting the potential impact of newly identified *MAGEL2* mutations based on their subcellular localization patterns, offering a novel angle for prognosis.

Subcellular localization significantly dictates the functional dynamics of proteins, establishing protein-protein interactions, post-translational modifications, and integration into cellular processes. Consequently, aberrant subcellular localization of proteins due to mutations has been acknowledged to trigger disease (12). As MAGEL2 is involved in multiple cellular signaling pathways including those related to retrograde endosomal transport (2), circadian rhythm (3), dendrite formation (13), or RNA metabolism (1), the nuclear localization of truncated MAGEL2 might interfere with these signaling cascades and lead to altered cellular responses. In addition to the disruption of normal cellular processes, the truncated protein might also acquire new functions in the nucleus, engaging in interactions with a different set of nuclear proteins compared to the WT. Truncation could yield abnormal protein products susceptible to aggregation, which may result in the aberrant regulation of gene expression and nuclear processes.

In conclusion, our study consistently shows a nuclear shift in truncated MAGEL2 variants, potentially disrupting cell functions and causing harmful effects associated with the SYS phenotype. Besides, we correlated the increased nuclear localization with the severity of AMC, providing valuable clinical implications. Collectively, our results shed new light on the complex molecular mechanisms underlying the pathogenicity of SYS-associated mutations and lay the groundwork for more research on the role of truncated MAGEL2 in the nucleus. Further studies could explore the mechanisms driving the nuclear localization of truncated MAGEL2, its nuclear functions, and therapeutic targeting to correct its import to the nucleus. Additionally, being aware that our study focused on exogenously expressed MAGEL2, it is essential to validate our results in the context of endogenous protein expression to establish their clinical relevance. Alternative approaches, such as CRISPR/Cas9-mediated tagging of endogenous MAGEL2, may enable more accurate assessment of its subcellular localization and potential interactions with other nuclear components.

## ACKNOWLEDGEMENTS

We would like to thank the patient’s association AESYS (Asociación Española del Síndrome Schaaf-Yang) for their generous financial support. We thank Monica Cozar for her technical assistance.

## AUTHOR CONTRIBUTIONS

RU, EA-C, AP-P and MC-P designed the cloning strategy. EA-C, MC-P and SG participated in the cloning of the plasmids. MC-P and SG conducted the cell culture, transfection, and immunofluorescence staining experiments. MC-P and JDG analyzed the ICC assays. MC-P and EA-C elaborated the corresponding figures and tables. SB, DG, RR and RU supervised the overall research process. MC-P, SB, RR, and RU drafted the manuscript. All authors have critically reviewed and approved the manuscript.

## ETHICAL APPROVAL

The study was approved by the Institutional Review Board (IRB00003099) of the Bioethical Commission of the University of Barcelona (5 October 2020) and Hospital Sant Joan de Déu (PIC-111-19).

