## Supplementary Information for "Subcellular localization of truncated MAGEL2 proteins: insight into the molecular pathology of Schaaf-Yang syndrome"

### SUPPLEMENTARY FIGURES

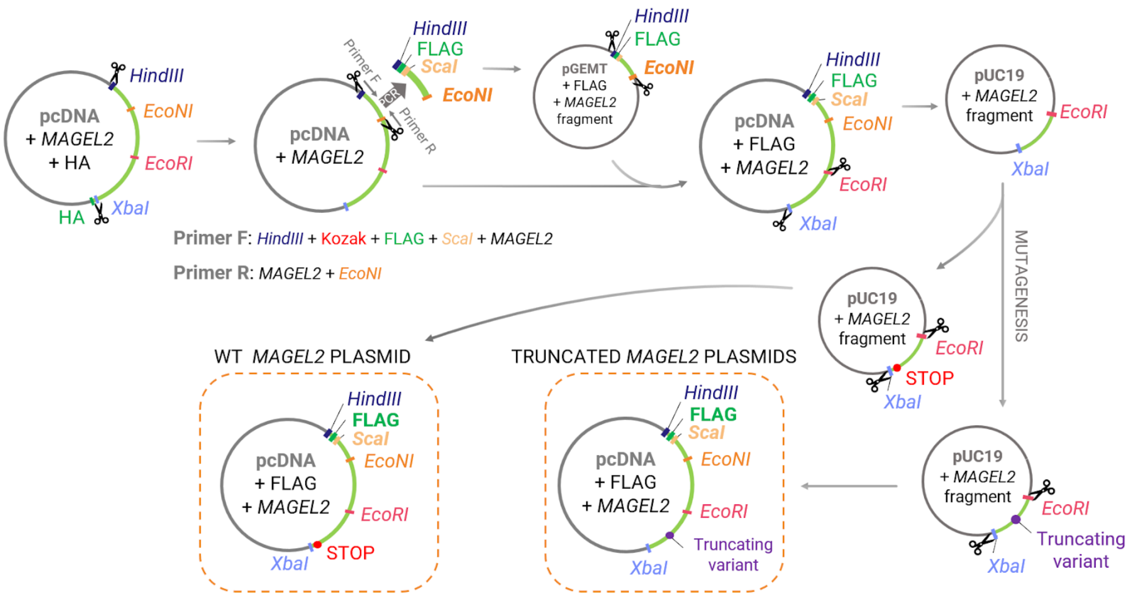


**Supplementary Figure 1**: Schematic representation of the cloning strategy used to generate the new FLAG-MAGEL2 plasmids from a previously described MAGEL2-HA vector.

### SUPPLEMENTARY MATERIALS AND METHODS

**Cloning**: The *MAGEL2* cDNA sequences of interest (ENST00000650528.1) were cloned in the mammalian expression vector pcDNA3.1(+) (Invitrogen, ThermoFisher Scientific) including a FLAG tag at the N-terminal end. To do so, the *MAGEL2* sequence was obtained from a previously described plasmid (1) using digestion enzymes *HindIII* (FD0504, ThermoFisher Scientific) and *XbaI* (FD0684, ThermoFisher Scientific). Then, 299 bps of the digested product were amplified by PCR using primers with an overhang to incorporate Kozak, FLAG and *ScaI* target sequences at N-terminal of *MAGEL2*. PCR primers are available in Supplementary Table 1. PCR product was cloned in a pGEM-T vector (A137A, Promega). Ligation was carried out using the Rapid DNA Dephos & Ligation Kit (Roche) following the manufacturer's instructions. Transformation of the ligation product was done in XL1-Blue (Agilent) competent cells with SOC media. Plasmids were purified (PureLinkTM Quick Plasmid Miniprep Kit, Invitrogen) and Sanger sequenced. Then, this FLAG-MAGEL2 N-terminal region was incorporated into the full pcDNA3.1(+) MAGEL2 plasmid using restriction enzymes *ScaI* (FD0434, ThermoFisher Scientific) and *HindIII* (FD0504, ThermoFisher Scientific), and the GFX PCR DNA and Gel Band Purification Kit (28903470, Cytiva).

To incorporate the six SYS patient mutations in the truncated FLAG-MAGEL2 plasmids, and a stop codon sequence in the WT plasmid, the last 1947 bps at C-terminal of *MAGEL2* were cloned in a pUC19 vector (SD1172, GenScript) using enzymes *EcoRI* (FD0274, ThermoFisher Scientific) and *XbaI* (FD0684, ThermoFisher Scientific). Mutations c.1850G>A, c.1912C>T, c.1996dupC, c.1996delC, c.2056_2066del, c.2118delT and a TAA stop codon sequence were introduced respectively by site-directed mutagenesis using the QuikChange Lightning Site-Directed Mutagenesis Kit (Agilent). Mutagenesis primers can be found in Supplementary Table 1.

**Cell culture**: HeLa cells were cultured in DMEM (Sigma-Aldrich, Merck) supplemented with 10% FBS (Gibco, LifeTechnologies) and 1% Penicillin–Streptomycin (Gibco, LifeTechnologies). The vectors were transfected into 60% confluent HeLa cells using LipofectamineTM 3000 (Invitrogen, ThermoFisher Scientific) and Opti-MEM (Gibco, LifeTechnologies).

**Immunocytochemistry**: HeLa cells were fixed in 4% Paraformaldehyde (PFA), permeabilized with 0,1 mol/L glycine and 0,1% Triton X-100 in PBS and blocked with 0,3 mol/L glycine, 0,05% Triton X-100 and 10% Normal Donkey Serum (#S30-100M, Merck Millipore) in PBS. Coverslips were incubated with anti-FLAG primary antibody (F3165, Sigma), Donkey anti-Mouse Cy2 antibody (#715-225-150, Jackson Immunoresearch), Phalloidin (#A30107, Invitrogen), and DAPI (#D1306, Invitrogen) and mounted with MOWIOL (#475904, Millipore). Images were acquired using a Zeiss confocal microscope LSM 880 and analysed with ImageJ (2).

### SUPPLEMENTARY TABLES

**Supplementary Table 1**: Primer sequences used in the cloning process of the FLAG-MAGEL2 plasmids.

| | | |
|---|---|---|
| c.1850G>A | Primer F | 5′-CTGGGCCTGCTAGGCCAGCGCCT-3′ |
| | Primer R | 5′-AGGCGCTGGCCTAGCAGGCCCAG-3′ |
| c.1912C>T | Primer F | 5′-GGGGGGAGCCTACCTCTGGGCCT-3′ |
| | Primer R | 5′-AGGCCCAGAGGTAGGCTCCCCCC-3′ |
| c.1996dupC | Primer F | 5′-CCTGGGCCTGCTGGGGGGGGTAGCT-3′ |
| | Primer R | 5′-AGCTACCCCCCCCAGCAGGCCCAGG-3′ |
| c.1996delC | Primer F | 5′-CCTGGGCCTGCTGGGGGGTAGCTG-3′ |
| | Primer R | 5′-CAGCTACCCCCCAGCAGGCCCAGG-3′ |
| c.2056_2066del | Primer F | 5′-CGCTCCAGCCTTCCGCCTGCAGTCTTGC-3′ |
| | Primer R | 5′-GCAAGACTGCAGGCGGAAGGCTGGAGCG-3′ |
| c.2118delT | Primer F | 5′-CGGTAGCAGCGGCAAATTTCCCCTGGGCT-3′ |
| | Primer R | 5′-AGCCCAGGGGAAATTTGCCGCTGCTACCG-3′ |
| TAA stop codon | Primer F | 5′-CCCCCTCCCCGCTAACTCGAGTCTAGAG-3′ |
| | Primer R | 5′-CTCTAGACTCGAGTTAGCGGGGAGGGGG-3′ |
| Kozak+FLAG+*ScaI* | Primer F | 5′-AAGCTTGCCACCATGGACTACAAAGACGATG ACGACAAGAGTACTTCGCAGCTAAGTAAGAAT-3′ |
| | Primer R | 5′-CCTAGGGCAGGAGGCTGGGT-3′ |
